# A glycoengineered antigen exploiting a conserved protein *O*-glycosylation pathway in the *Burkholderia* genus for diagnosis of glanders infections

**DOI:** 10.1101/2020.08.27.271338

**Authors:** Guanbo Wang, Lena Glaser, Nichollas E. Scott, Yasmine Fathy Mohamed, Rebecca Ingram, Karine Laroucau, Miguel A. Valvano

## Abstract

We recently described a protein *O*-glycosylation pathway conserved in all species of the *Burkholderia* genus that results in synthesis and incorporation of a trisaccharide glycan to membrane-exported proteins. Here, we exploited this system to construct and evaluate a diagnostic tool for glanders. *Burkholderia mallei*, the causative agent of glanders, is a highly infectious and fatal zoonotic pathogen that mainly infects horses, mules, donkeys and occasionally humans. A highly sensitive and specific diagnostic tool is crucial for the control, elimination and eradication of *B. mallei* infections. We constructed plasmids carrying synthetic genes encoding a modified, previously unannotated *Burkholderia* glycoprotein containing three glycosylation sequons fused to the cholera toxin B-subunit. The resulting proteins were glycosylated in the *B. cenocepacia* K56-2 parental strain, but not in glycosylation-deficient mutants, as determined by SDS-PAGE and fluorescent lectin blots. One of these glycoproteins was used as an antigen in ELISA and western blots to screen a panel of serum samples collected from glanders-infected and healthy horses previously investigated by complement fixation test and indirect ELISA based on a semi-purified fraction of *B. mallei*. We show that ELISA and western blot assays based on our glycoprotein antigen provide 100 % specificity, with a sensitivity greater than 88%. The glycoprotein antigen was recognized by serum samples collected from patients infected with *B. pseudomallei, B. mallei, B. multivorans* and *B. cenocepacia*. Our results indicate that protein *O*-glycosylation in *Burkholderia* can be exploited as a biomarker for diagnosis of *Burkholderia*-associated infections.

**IMPORTANCE:** Glanders is a severe zoonotic disease caused by the Gram-negative bacterium *Burkholderia mallei*, which affects horses, mules and donkeys, as well as humans. *B. mallei* is also considered a category B biothreat agent. Due to insufficient pathognomonic symptoms in the early stages of glanders, diagnosis can be difficult. Complement fixation is the most accurate and reliable serological test prescribed by the World Organization for Animal Health; however, this test has a considerable number of false-positive results. We have recently described a conserved protein *O*-glycosylation pathway present in all species of the *Burkholderia* genus; we also demonstrated that *Burkholderia-*infected humans develop anti-glycan antibodies. Here, we exploited this system to construct and evaluate a synthetic glycoengineered protein antigen as a diagnostic tool for glanders. Our results show 100 % specificity in the detection of antibodies from infected horses, indicating that protein *O*-glycosylation in *Burkholderia* can be exploited as a biomarker for diagnosis of *Burkholderia*-associated infections.

## INTRODUCTION

Glanders is a severe zoonotic disease caused by the Gram-negative bacterium *Burkholderia mallei*. Horse, mules and donkeys are natural hosts, while transmission from animals to humans has occurred in people who are working closely with infected animals (1). The signs and symptoms of infection can be either acute or chronic depending on the individual. Acute infections occur frequently in donkeys and mules; they are usually fatal without treatment, showing signs of septicemia, high fever, weight loss and compromise of the respiratory system. However, glanders generally manifests with a more chronic course in horses (2). Although the disease has been successfully eradicated in North America, Australia and Europe, there are reports of glanders in Asia, the Middle East and South America, and hence it is considered as a re-emerging disease (3-7). Also, *B. mallei* is considered a category B biothreat agent (8).

Due to insufficient pathognomonic symptoms, especially in the early stages of glanders, diagnosis can be difficult. Direct isolation and molecular identification of *B. mallei* from infected tissues, cutaneous lesions and nasal exudates is the most accurate way to confirm the infection. However, this is generally limited by poor sensitivity due to low bacteria load, as subclinical and latent infections are commonly manifested in horses (2). Mallein and serological tests are frequently used for diagnosis of glanders in endemic areas (9). The mallein test is a hypersensitive skin test based on a water-soluble protein extracted from the microorganism, but it is not recommended by the World Organization for Animal Health (OIE) due to animal welfare concerns since the mallein antigen needs to be injected into the horse (http://www.oie.int/fileadmin/Home/eng/Health_standards/tahm/3.05.11_GLANDERS.pdf.). Complement fixation test (CFT) is the most accurate and reliable serological test prescribed by the OIE for international trade of equines. Although this test is believed to have a high specificity, a considerable number of false-positive results were observed when different antigens and assay protocols were applied (10, 11); complexity, anti-complementary reactions and poor standardization are other shortcomings of the CFT (12). The Rose Bengal plate agglutination test is a rapid agglutination assay using a heat-inactivated bacterial suspension colored with Rose Bengal; it is fast and simple but requires antigens purified from *B. mallei* cultures (9, 13).

Recently, several purified protein antigens of *B. mallei* have been tested in ELISA and western blot showing promising results for diagnosis of glanders. A western blot technique based on a partially-purified lipopolysaccharide-containing antigen shows higher specificity compared to CFT (14). Moreover, indirect ELISAs based on the intracellular motility A protein (BimA) (15, 16), the type 6 secreted TssB (17) and Hcp1 (18) proteins, the heat shock protein GroEL (19) and a semi-purified fraction of *B. mallei* have been developed (11). However, these methods require more investigation using large-scale sample surveys and optimization to improve specificity and sensitivity (18). Previous work has identified a *O*-glycosylation (*ogc*) gene cluster in *B. cenocepacia* encoding enzymes for the synthesis of a lipid-linked trisaccharide, which consists of a β-Gal-(1,3)-α-GalNAc-(1,3)-β-GalNAc (20) and is incorporated in membrane-exported *Burkholderia* proteins by the PglL *O-*glycosyltransferase (*O-*Tase) (21). We also found this pathway is conserved in all members of the *Burkholderia* genus including *B. mallei* (20). More imporantly, we detected glycan-specific antibodies in sera from patients infected with various types of *Burkholderia* infections including glanders, indicating that natural infection elicits a humoral response against the *Burkholderia*-specific glycan (20). In this study, we describe the construction of a glycoengineered antigen that can be exploited as a highly specific diagnostic tool for glanders, and potentially other *Burkholderia* related infections.

## MATERIALS AND METHODS

### Bacteria strains and growth conditions

*Escherichia coli* K-12 strain DH5α was used for cloning experiments to construct the appropriate plasmids. *B. cenocepacia* K56-2 strain and the glycosylation-deficient mutants Δ*pglL* and Δ*ogc* (20) were used to produce glycosylated and unglycosylated proteins, respectively. Bacterial strains were grown at 37°C in Luria-Bertani (LB) medium supplemented with appropriate antibiotics. For *B. cenocepacia* K56-2 and mutants carrying the plasmid pDA12, bacteria were grown with 100 μg ml^-1^ tetracycline. *E. coli* DH5α carrying the plasmid pDA12 was grown with 30 μg ml^-1^ tetracycline. *E. coli* DH5α carrying the plasmid pRK2013 was grown with 40 μg ml^-1^ kanamycin.

### Plasmid constructions

To obtain high-level expression of recombinant proteins in *B. cenocepacia* we used the constitutive expression vector pDA12 (22). The chimeric genes encoding the respective variations of recombinant proteins, BCAL2737a, CtxB-BCAL2737a, BCAL2737a-CtxB were synthesized (Eurofins Genomics) and gene fragments subcloned into pDA12 to generate recombinant plasmids pDA12-BCAL2737a, pDA12-BCAL2737a-CtxB, pDA12-CtxB-BCAL2737a. Positive clones were screened by colony PCR using primers sets: 5’-ACTCTCGCATGGGGAGACCC-3’ and 5’-TTTGATGTTATGGAGCAGCAACGAT-3’. The Go Taq® DNA polymerase (Promega, UK) was used for PCR amplification, conditions were 4 min at 95°C, 34 cycles of 95°C for 30 s, 58°C for 40 s, and 72°C for 3 min, and a final extension of 7 min at 72°C.

The resulting recombinant plasmids were mobilized from *E. coli* DH5α into *B. cenocepacia* K56-2 and the isogenic mutants Δ*pglL* and Δ*ogc* by triparental mating using an *E. coli* DH5α carrying the helper plasmid pRK2013 (23). Exconjugants were selected for on LB agar plates with 50 μg ml^-1^ gentamicin, 200 μg ml^-1^ ampicillin and 120 μg ml^-1^ tetracycline. Successful exconjugants were identified by colony PCR as described above.

### Expression and purification of recombinant proteins

For expression, bacteria were grown in 1 liter of LB medium with 100 μg ml^-1^ tetracycline overnight at 37°C. For purification, cells were centrifuged for 20 min at 3,220 *×g* followed by two washes with PBS at 4°C. The pellet was resuspended in 20 ml Tris-HCl buffer (100 mM, pH 8.0). DNAse (5 mg ml ^-1^), lysozyme (1 mg ml ^-1^), and protease inhibitors (1 tablet, Roche) were added and the mixture was incubated for 15 min. Cells were lysed using a cell disrupter (Constant systems Ltd., Northants, UK) at two cycles of 30 kPsi. The lysed cells were centrifuged for 20 min at 17,696 *×g* at 4°C and the resulting supernatant was immediately filtered using a 0.2 μm membrane (Fisher Scientific, UK). The filtrate was applied to a ÄKTA protein purification system (GE Healthcare Life Sciences), which was equipped with a His-trap column. Proteins were eluted using a linear gradient of up to 100% elution buffer (100 mM Tris-HCl [pH 8.0], 500 mM NaCl, 500 mM Imidazole, glycerol 2%). The collected fractions were analyzed by 12% SDS-PAGE gels and western blot.

### Western blot analysis of protein glycosylation and equine serum samples

The presence of the trisaccharide glycan in purified proteins was analyzed by western blot using biotinylated peanut agglutinin (PNA), a lectin that recognizes the terminal disaccharide of the *Burkholderia* glycosylation trisaccharide (20, 24). Purified proteins were run on 12% SDS-PAGE gels and then blotted onto nitrocellulose membranes using the BIO-RAD Trans-Blot® Turbo™ Transfer System. Membranes were blocked overnight in casein solution (Sigma-Aldrich) diluted in Tris-buffered saline, 0.1% Tween 20 (TBST). Mouse anti-His antibody (1:12000) (Sigma-Aldrich) and biotinylated PNA (1:6000) (Vector Laboratories) were applied as primary antibodies. The mouse anti-His antibody was added first and incubated at room temperature for 1 h. Biotinylated PNA was added to this solution followed by incubation for another 30 min. After four washes with TBST, IRDye® 680RD goat anti-mouse antibody (1:12000) (Li-Cor, Biosciences) and IRDye® 800CW Streptavidin (1:12000) (Li-Cor, Biosciences) were applied as secondary antibodies. The goat anti-mouse antibody was applied first and incubated at room temperature for 30 min. Streptavidin was added to the solution and further incubated for 20 min. After four washes, as before, the membranes were visualized using the Odyssey Infrared Imaging System (Li-Cor, Biosciences).

To detect antibodies in equine serum samples, an optimized western blot protocol was established. Briefly, the same protocol was used for preparing nitrocellulose membranes with purified proteins. The membranes were cut into regular strips and each strip incubated in a final volume of 2 ml of each serum sample, diluted 1:1000 in TBST for 1 h at room temperature. After four washes with TBST, the membranes were then incubated with biotinylated goat anti-horse IgG antibody (1:12000) (Abcam, UK). After four more washes, the membranes were stained with IRDye® 800CW Streptavidin for 30 min and visualized in the Odyssey Infrared Imaging System.

### ELISA to detect antibodies toward glycan in equine serum samples

An optimized ELISA protocol was established after several trials to determine the optimal antigen concentration and serum dilutions. To reduce the background, serum samples were adsorbed with unglycosylated protein by incubating the samples with the unglycosylated CtxB-BCAL2737a at a concentration of 1 μg ml^-1^ for 2 h at 37°C and overnight at 4°C in a 96-well Nunc MaxiSorp plate. Serum samples were then collected and used for detection by ELISA using the following procedures: 96-well Nunc MaxiSorp plates were coated with 50 μl of purified glycosylated and unglycosylated proteins diluted in coating buffer (carbonate/bicarbonate, 100 mM, pH 9.6) to reach a final concentration of 31 pg ml^-1^. The plates were covered by plastic adhesive film and incubated at 4°C overnight. Coating solution was removed, and plates were washed with 300 μl phosphate buffer saline (PBS) containing 0.05% Tween 20 (PBST). Additional blocking was achieved by adding 300 μl of blocking buffer (BSA 5% in PBS). Plates were covered and incubated at room temperature for 1 h and then washed 3 times with PBST. Fifty μl of each serum sample, diluted in half-strength blocking buffer, was added to the wells and incubated for 90 min at room temperature. Biotinylated PNA and PBST were used as a positive and negative controls. Plates were washed 4 times with PBST. Fifty μl of biotinylated rabbit anti-horse IgG secondary antibody (Abcam, UK) diluted to 1:5000 was added and incubated for 1 h at room temperature. Plates were washed 5 times with PBST. Fifty μl of Streptavidin-Horseradish Peroxidase diluted 1:200 in PBS was added to each well and incubated for 1 h in the dark at room temperature. After washing 4 times with PBST, 50 μl of the substrate solution 3,3’,5,5’-tetramethylbenzidine was added per well and incubated in the dark at room temperature for 10 min. After enough color development, 30 μl of stop solution (2N H_2_SO_4_) was added and the absorbance of each well was read with POLARstar Omega microplate reader (BMG LABTECH, Ortenberg, Germany) at 450 nm.

### Horse serum samples

Thirty-two serum samples from horses naturally and experimentally infected with *B. mallei*, and healthy controls were used for analysis. Positive samples were characterized by the complement fixation test (CFT). An indirect ELISA based on a semi-purified fraction of *B. mallei* described previously (18) was carried out for comparison. The CFT was performed according to the OIE manual. The indirect ELISA was performed according to the manufacturer’s instructions (IDvet, Grabels, France). Optical density (OD) were read at 450 nm and results were expressed according to the manufacturer’s instructions as: S/P = (OD_Sample-OD_Negative control)/ (OD_Positive control-OD_Negative control) x 100. S/P values over 40% were considered positive.

### Human serum samples

Nine serum samples from *Burkholderia*-infected patients that were previously reported to interact with *O*-glycan (20) were selected for western blot analysis performed as described above. Among these samples, two are from *B. multivorans*-infected patients, three samples are from *B. cenocepacia*-infected patients, one sample is from a *B. mallei*-infected patient and three samples are from *B. pseudomallei*-infected patients.

### Proteomic analysis

Gel separated CtxB-BCAL2737a samples were excised and destained in destaining solution (100 mM NH_4_HCO_3_, 50% ethanol (EtOH), pH 8.0) for 20 min at room temperature. with shaking (750 rpm). Destained gel samples were then washed with 100% EtOH for 10 min at room temperature and dried by vacuum centrifugation for 20 min. Dried gel samples were rehydrated with 10 mM dithiothreitol in 100 mM NH_4_HCO_3_ and disulfide reduction carried out for 60 min at 56°C with shaking. The reducing buffer was removed, and the gel samples washed twice in 100% EtOH for 10 min to remove residual dithiothreitol. Samples were alkylated with 55 mM iodoacetamide in 50 mM NH_4_HCO_3_ in the dark for 45 min at room temperature. Alkylated samples were washed with 100 mM of NH_4_HCO_3_ for 10 min at room temperature with shaking and then dehydrated with 100% EtOH before being vacuum centrifugation for 20 min. Dried reduced/alkylated samples were then rehydrated with 20 ng/µl trypsin in 100 mM NH_4_HCO_3_ (Promega, Madison WI) on ice for 1 h. Excess trypsin solution was removed, gel pieces covered in 40 mM NH_4_HCO_3_ and then incubated overnight at 37°C. The supernatant, containing peptides of interest, were concentrated and desalted using C_18_ stage tips (25, 26) before analysis by liquid chromatography-mass spectrometry.

Desalted tryptic peptides were resuspended in Buffer A* (0.1% trifluoroacetic acid, 2% Acetonitrile) and separated using a two-column chromatography set up composed of a PepMap100C18 20mm x 75μm trap and a PepMap C18 500 mm × 75 μm analytical columns (Thermo Scientific, San Jose CA). Samples were concentrated onto the trap column at 5 μl min^-1^ for 5 min with Buffer A (0.1% formic acid, 2% Acetonitrile) and infused into an LTQ-Orbitrap Elite (Thermo Scientific, San Jose CA) at 300 nl min^-1^ via the analytical column using a Dionex Ultimate 3000UPLC (Thermo Scientific). A 90-min gradient was run from 2% Buffer B (0.1% formic acid, 80% Acetonitrile) to 32% B over 51 min, then from 32% B to 40% B in the next 5 min, then increased to 100% B over 2 min period, held at 100% B for 2.5 min, and then dropped to 0% B for another 20 min. The LTQ-Orbitrap Elite was operated in a data-dependent mode automatically switching between mass spectrometry (MS), collision-induced dissociation (CID), electron transfer dissociation (ETD), and higher energy collision-induced dissociation (HCD) MS-MS.

Identification of glycopeptides in CtxB-BCAL2737a was accomplished using MaxQuant (v1.5.3.30) (27). Searches were performed against the predicted amino acid sequence of CtxB-BCAL2737a with carbamidomethylated cysteine set as a fixed modification. Searches were performed with trypsin allowing two missed cleavage events and the variable modifications of oxidation of methionine, the *Burkholderia* glycans (Glycan 1; 568.211 Da elemental composition: C22H36O15N2 and Glycan 2; 668.227 Da elemental composition C26H40O18N2) and acetylation of protein N-termini. The precursor mass tolerance was set to 20 parts-per-million (ppm) for the first search and 10 ppm for main search, with a maximum false discovery rate of 1.0% set for protein and peptide identifications. The resulting outputs were processed within the Perseus (v1.4.0.6) (28) analysis environment to remove reverse matches and common proteins contaminates prior to further analysis. Glycopeptide identified were manually assessed according to the guidelines of Chen *et al*. (29), annotated manually with the aid of Protein Prospector tool MS-Product (http://prospector.ucsf.edu/prospector/cgi-bin/msform.cgi?form=msproduct). The MS proteomics data were deposited in the ProteomeXchange Consortium via the PRIDE partner repository (30, 31) with the dataset identifier PXD018788. Data can be accessed under username, password: C7kqIWps.

### Analysis of BCAL2737a degradation in *Burkholderia* strains

As BCAL2737a was not annotated in the reference *B. cenocepacia* strain J2315 (see Results), we re-analyzed the previously published *B. cenocepacia* K56-2 peptidome datasets (Pride accession: PXD014614) (32) including BCAL2737a. Searches were performed against the predicted amino acid sequence of BCAL2737a and the K56-2Valvano proteome (NCBI: Taxonomy ID: 985076) with identical setting as above. The resulting outputs were processed in the Perseus (v1.5.0.9) analysis environment to remove reverse matches and common protein contaminates prior to further analysis. Peptides identified across samples were visualized in heat maps using R (https://www.r-project.org/).

### Statistical analysis

Means and standard deviations were calculated using GraphPad Prism 8 software (GraphPad Prism 8; GraphPad Software, USA). Three independent repeats were tested for every sample in ELISA. The cutoff value is calculated based on OD450 nm value of samples tested negative in CFT (excluding inconclusive results). Samples were considered positive when the absorbance reading exceeds the cutoff value computed by two standard deviations above the mean of the negative control. The specificity and sensitivity of the ELISA and western blot tests were determined using the CFT as a standard according to the following formulas:

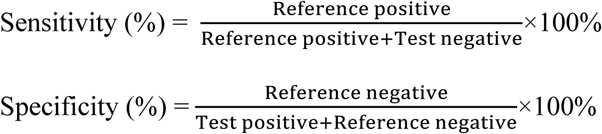

Inconclusive CFT results were not considered for these calculations.

## RESULTS

### Identification of a protein-glycan glycoconjugate antigen for antibody detection

In a previous study characterizing the general protein glycosylation system in *B. cenocepacia*, one of the most abundant and highly glycosylated glycopeptides found, designated A0K9U9, could not be ascribed to any annotated protein in the database of the type strain J2315 (21). To localize the gene encoding this peptide, its 71 amino acid sequence was used as query in Artemis (33) to search the translations in all frames for *B. cenocepacia* type strain J2315. This resulted in the identification of a short open reading frame in chromosome 1 (at base pairs 3008284-3008496) located in the intergenic region between BCAL2737 (encoding a putative pseudouridine synthase) and BCAL2738 (encoding a putative exported protein), which was preceded by a ribosomal binding site sequence (Fig. 1); we designated this open reading frame BCAL2737a. The predicted BCAL2737a gene encoded a 71-amino acid polypeptide; it also contained a 21-amino acid signal peptide, predicted by SignalP v.5.0 (34), as expected for a glycosylated protein since glycosylation takes place in the periplasm. The BCAL2737a protein did not show any homology with known proteins in the database. The mature polypeptide contained 3 serine residues (Fig. 1) that become *O*-glycosylated based on experimental evidence by mass spectrometry (21). Therefore, the BCAL2737a polypeptide was selected for further analysis since because of its small mass, combined with 3 glycosylation sites, it could be an appropriate antigen to detect antibodies against *Burkholderia O-*glycans.

**FIG 1.**
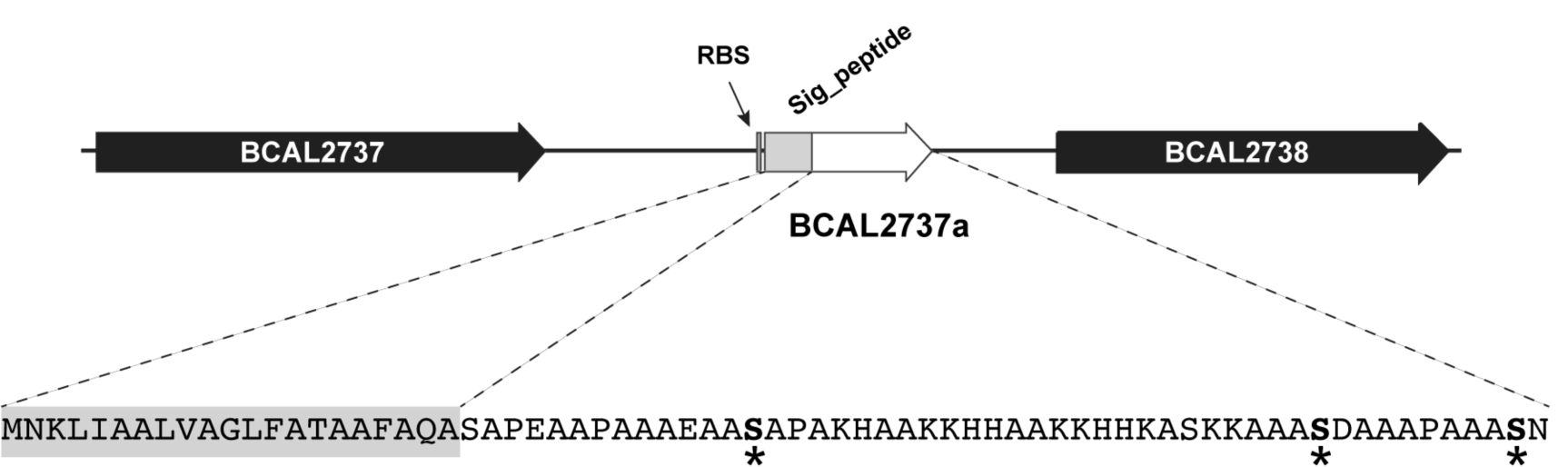
Localization of BCAL2737a in the large chromosome of *B. cenocepacia* J2315. BCAL2737a, encoding a short polypeptide of unknown function, was identified in the intergenic region between BCAL2737 (encoding a putative pseudouridine synthase) and BCAL2738 (encoding a putative exported protein. The gene is preceded by a Shine-Dalgarno sequence (RBS). The BCAL2737a polypeptide has a signal peptide (Sig_peptide) as predicted by SignalP v.5.0, resulting in a 50-amino acid mature protein; the *O*-glycosylation sites, experimentally determined by mass spectrometry (21), are indicated by asterisks.

### Unglycosylated BCAL2737a can be stabilized as a protein fusion with the cholera toxin B subunit (CtxB)

BCAL2737a was expressed as a 10x-His tag C-terminal fusion (Fig. 2A) and examined for glycosylation using PNA, and for protein expression using an anti-His monoclonal antibody. This experiment indicated that BCAL2737a can be detected by both methods when expressed in the parental *B. cenocepacia* K56-2 (Fig. 2B). The lectin blots showed also several non-specific bands attributable to endogenous biotinylated bacterial proteins in the lysate, which were detected by the secondary fluorescent streptavidin (Fig. 2C, and asterisks in Fig. 2B and D). However, the BCAL2737a was undetectable in both the Δ*ogc* and Δ*pglL* mutants, which lack the biosynthetic cluster for the glycan assembly and the PglL *O-*Tase, respectively (Fig. 2B and D). This was not unexpected since a subset of the *Burkholderia* glycoproteome is proteolytically degraded (32). Therefore, as a strategy to stabilize BCAL2737a, we also generated versions of this proteins that were N- and C-terminally fused with the cholera toxin B subunit (CtxB) and contained a C-terminal 10x-His tag (Fig. 2A). Both proteins were expressed and glycosylated in K56-2, but only the CtxB-BCAL2737a fusion protein remained stable in the glycosylation-defective mutants (Fig. 2B and D). These results suggest that in the absence of glycosylation BCAL2737a can only be stabilized when it is C-terminally fused to CtxB. Therefore, we continued our studies with this fusion protein.

**FIG 2.**
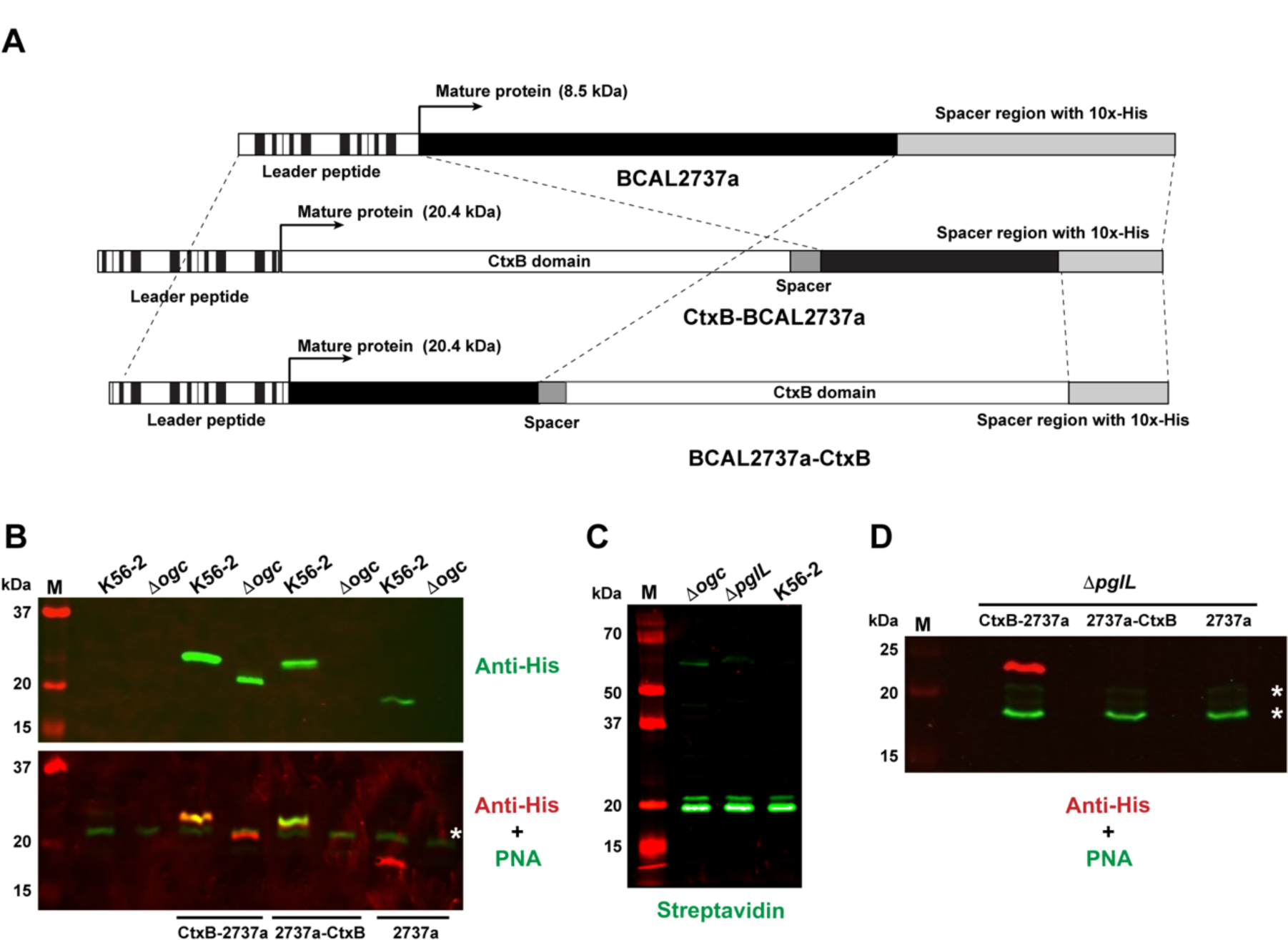
Protein constructs containing BCAL2737a and western blots demonstrating the effects of a lack of glycosylation in Δ*ogc* and Δ*pglL* strains. (A) Map of the CtxB-BCAL2737a, BCAL2737a-CtxB and BCAL2737a recombinant polypeptides indicating their functional domains. Maps are not drawn to scale. (B) Comparison of protein expression and glycosylation in K56-2 and Δ*ogc* strains with no plasmid and with plasmids expressing CtxB-BCAL2737a (CtxB-2737a), BCAL2737a-CtxB (2737a-CtxB), and BCAL2737 (2737a). (C) Detection of endogenously biotinylated proteins by direct application of fluorescent streptavidin to the western blot. Lanes M, molecular mass markers, kDa, kilodaltons. (D) Protein expression and glycosylation in Δ*pglL* using the same plasmids as in panel B. Asterisks in panels B and D, indicate the most abundant non-specific, endogenously biotinylated proteins

To confirm that CtxB-BCAL2737a is efficiently glycosylated, the protein was purified by Ni^2+^-affinity chromatography from K56-2 and from Δ*pglL* as a control. Western blot analysis revealed that purified CtxB-BCAL2737a expressed in Δ*pglL* exhibited decreased electrophoretic mobility, suggesting it corresponded to an unglycosylated form (Fig. 3A), in agreement with the results found with crude lysates (Fig. 2B and D). Similarly, only the protein expressed in K56-2 was detected with PNA, indicating the correct trisaccharide glycan residues were present. Glycosylation was also verified by mass spectrometry confirming at least three serine residues within BCAL2737a are glycosylated (Fig. 3B, Supplementary Dataset). To further support the degradation of unglycosylated BCAL2737a peptidome datasets from Oppy et al. (32) were reanalyzed against a K56-2 proteome containing the BCAL2737a protein sequence. The new analysis identified multiple peptide breakdown products previously not assigned to known proteins, which were more abundant and intense in the glycosylation null strain Δ*pglL* compared to K56-2 and the *pglL* complemented strain (Fig. 3C). Together, the results of these experiments demonstrate that CtxB-BCAL2737a is glycosylated at three sites, as the native protein and BCAL2737a is proteolytically degraded in the absence of glycosylation.

**FIG 3.**
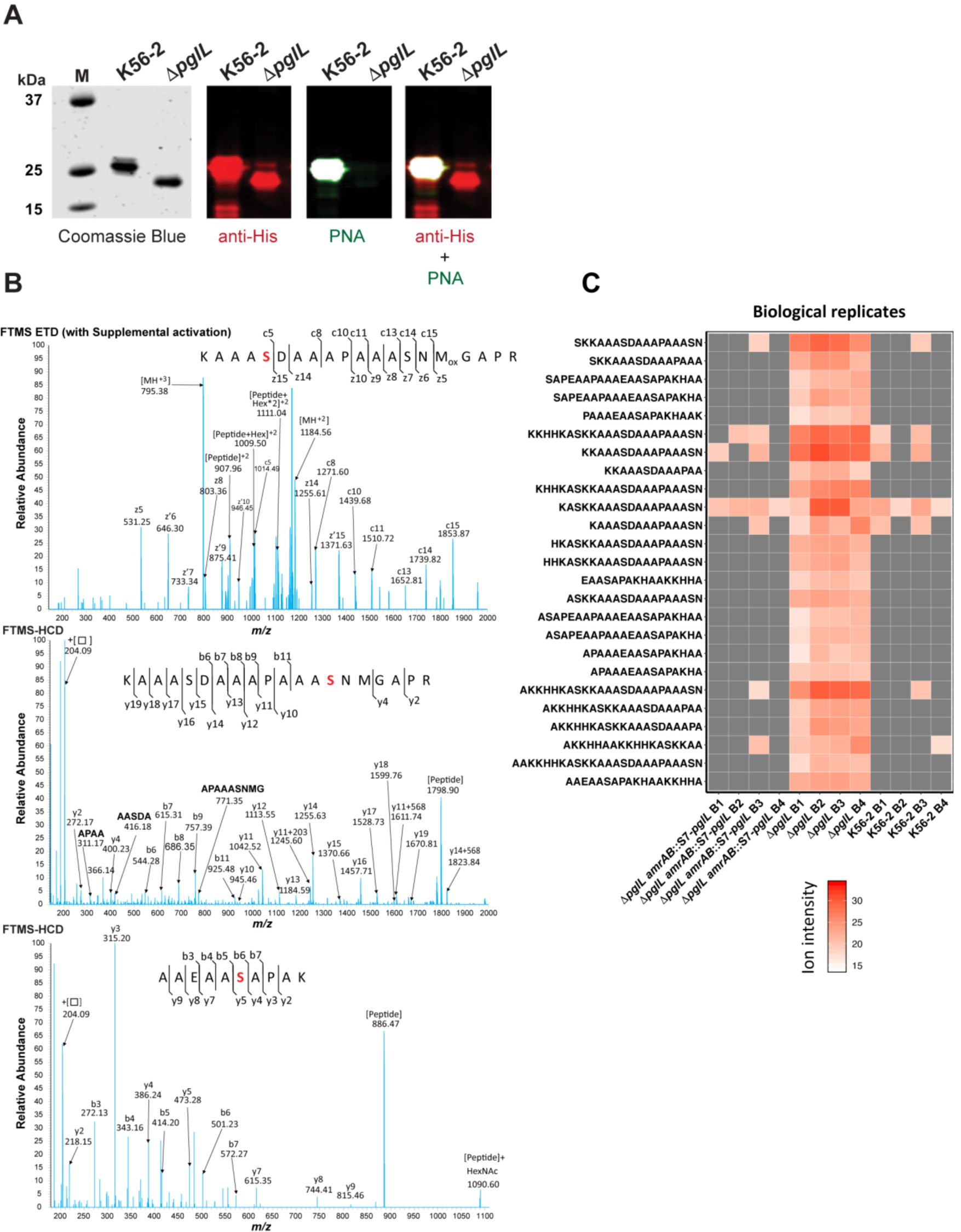
Characterization of purified CtxB-BCAL2737a. (A) CtxB-BCAL2737a was purified from strains K56-2 and Δ*pglL* by Ni^2+^-affinity chromatography and analyzed by SDS-PAGE (Coomassie Blue) and by western blot with anti-His and PNA, as indicated. M, molecular mass standards; kDa, kilodaltons. (B) The band corresponding to glycosylated BCAL2737a (purified from strain K56-2) was extracted from the gel, processed as indicated in Material and Methods, and the glycopeptides analyzed by MS-MS in an Orbitrap Fourier transform mass spectrometer (FTMS) with electron transfer dissociation (ETD) and higher energy collision-induced dissociation (HCD). The data confirm all three serine residues previously reported (21), also indicated in Fig. 1, are modified in CtxB-BCAL2737a. The modified serine residues are shown in red. (C) Heatmap of BCAL2737a peptides observed in peptidome preparations of *B. cenocepacia* K56-2. Peptidome datasets from Oppy et al. (32) were reanalyzed against a K56-2 proteome containing the BCAL2737a protein sequence, resulting in the identification of multiple peptide breakdown products previously not assigned to known proteins, which are clearly both more numerous and more intense in the glycosylation null strain Δ*pglL* compared to K56-2 and the *pglL* complemented strain (32). B, Biological replicate.

### Glanders-infected horse serum samples contain anti-*O*-glycan antibodies

An indirect ELISA was employed to evaluate if glycosylated CtxB-BCAL2737a can be used for diagnosis of glanders. Serum samples from thirty-two horses diagnosed with glanders and healthy controls, previously tested by CFT and indirect ELISA based on semi-purified *B. mallei*, were examined for antibodies against the glycosylated CtxB-BCAL2737a protein (Table 1). From these, 12 samples testing positive in both CFT and *B. mallei* indirect ELISA also gave positive results in the ELISA based on glycosylated CtxB-BCAL2737a. They included 7 samples (14-566_1831, 14-566_1850, 14-566_1861, 14-566_1899, 14-566_1917, 16-2439_81, 16-2439_85) from South America (all from confirmed cases of glanders at different locations), and 5 samples (19-5577_3, 19-5577_4, 19-5577_6, 19-5577_37, 19-5577_38) from the Middle East (from an outbreak in the same farm). However, three of the positive samples (19-5577_3, 19-5577_6, 16-2439_81) showed non-specific interactions with the unglycosylated control, as indicated by slightly higher OD values than the cut-off value (Fig. 4). The remaining positive samples had strong interaction with glycosylated protein only, indicating these serum samples contained specific anti-glycan antibodies. Further, two samples (18-1575/2905, 19-5577_39) gave inconclusive results by CFT, but these two samples test negative in the indirect ELISA based on semi-purified *B. mallei* (Table 1). Based on the negative results by the ELISA using glycosylated CtxB-BCAL2737a, we considered these samples as negative. Interestingly, three samples (Horse_1_(Océane)_ D_19/04_, Horse_2_(Poupée)_ D_19/04_, and Horse_3_(Princese)_ D_26/04_) from an immunization trial with heat-inactivated *B. mallei* tested negative in ELISA based on glycosylated CtxB-BCAL2737a as opposed to the results in CFT and ELISA based on semi-purified *B. mallei* (Table 1). From these results, we concluded that the ELISA based on glycosylated CtxB-BCAL2737a was accurate with sensitivity and specificity of 88.23% and 100%, respectively (Table 2).

**TABLE 1.** Comparison of antibody detection in 32 horse serum samples by CFT, indirect ELISA based on semi-purified *B. mallei*, and assays based on glycosylated CtxB-BCAL2737a

| Sample | Origin | CFT<sup>a</sup> | iELISA<sup>b</sup>-<br>semi-purified<br>*B. mallei*<br>(S/P%) | iELISA<sup>c</sup><br>CtxB-BCAL2737a | Western blot<br>CtxB-BCAL2737a |
| --- | --- | --- | --- | --- | --- |
| 18-117/302 | France | - | - (3%) | - | - |
| 18-1575/2905 | France | +/- (1) | - (4%) | - | - |
| 13-3290/7936 | Ireland | - | - (3%) | - | - |
| 13-3290/7937 | Ireland | - | - (1%) | - | - |
| 13-3290/7938 | Ireland | - | - (0%) | - | - |
| 13-3290/7939 | Ireland | - | - (12%) | - | - |
| 13-3290/7940 | Ireland | - | - (1%) | - | - |
| 18-103/214 | Tunisia | - | - (7%) | - | - |
| 18-103/215 | Tunisia | - | - (3%) | - | - |
| 18-103/216 | Tunisia | - | - (4%) | - | - |
| 18-103/218 | Tunisia | - | - (7%) | - | - |
| 14-566_1831 | South America | + (421) | + (149%) | + | + |
| 14-566_1850 | South America | + (44431) | + (128%) | + | + |
| 14-566_1861 | South America | + (42) | + (180%) | + | + |
| 14-566_1899 | South America | + (4) | + (134%) | + | + |
| 14-566_1917 | South America | + (432) | + (173%) | + | + |
| 16-2439_81 | South America | + (44442) | + (63%) | + | + |
| 16-2439_85 | South America | + (4442) | + (146%) | + | + |
| 19-5577_3 | Middle East | + (444443) | + (73%) | + | - |
| 19-5577_4 | Middle East | + (4442) | + (107%) | + | + |
| 19-5577_6 | Middle East | + (444441) | + (119%) | + | + |
| 19-5577_37 | Middle East | + (4441) | + (96%) | + | + |
| 19-5577_38 | Middle East | + (333) | + (47%) | + | - |
| 19-5577_39 | Middle East | +/- (1) | - (29%) | - | - |
| Horse_1 (Océane) | D08/02 immunization trial<sup>d</sup> | - | - | - | - |
| Horse_2 (Poupée) | D08/02 immunization trial<sup>d</sup> | - | - | - | - |
| Horse_3 (Princesse) | D08/02 immunization trial<sup>d</sup> | - | - | - | - |
| Horse_4 (Quirina) | D08/02 immunization trial<sup>d</sup> | - | - | - | - |
| Horse_1 (Océane) | D19/04 immunization trial<sup>d</sup> | + (32) | + (47%) | - | - |
| Horse_2 (Poupée) | D19/04 immunization trial<sup>d</sup> | + (444) | + (60%) | - | + |
| Horse_3 (Princesse) | D26/04 immunization trial<sup>d</sup> | + (443) | + (119%) | - | + |
| MRI#1 | lyophilized serum from a naturally infected horse | - | + (111%) | - | - |

<sup>a</sup> Numbers in parenthesis indicate the magnitude of the CFT reported from the combination of two parameters: the last dilution giving a positive result combined with the intensity of hemolysis inhibition indicated as 0 = 0%, 1 = 25%, 2 = 50%, 3 = 75%, and 4 = 100%. Typically, sera were tested at dilutions 1/5, 1/10, 1/20, 1/40 and 1/80 and in some strong positives at 1/160.
<sup>b</sup> Numbers in parenthesis result from the following formula: (OD\_Sample-OD\_Negative control)/(OD\_Positive control-OD\_Negative control) x 100. Values above 40% are considered positive.
<sup>c</sup> based on the cutoff OD value indicated in Fig. 4.
<sup>d</sup> These sera are from an immunization trial with a heat-inactivated *B. mallei* suspension performed in three horses; D, day/month) of sampling. 08/02: day 0; 19/04 and 26/04, days 63 and 70 post-immunization.

**TABLE 2.** Results of ELISA and western blot analysis of 32 horse serum samples

| Assay (Results)<sup>a</sup> | Reference test<sup>b</sup> (+) | Reference test<sup>b</sup> (-) |
| --- | --- | --- |
| ELISA (+) | 15 | 0 |
| ELISA (-) | 2 | 15 |
| Western blot (+) | 15 | 0 |
| Western blot (-) | 2 | 15 |

<sup>a</sup> 2 samples giving inconclusive results (+/-) in CFT were excluded from these comparisons.
<sup>b</sup> CFT was used as reference

**FIG 4.**
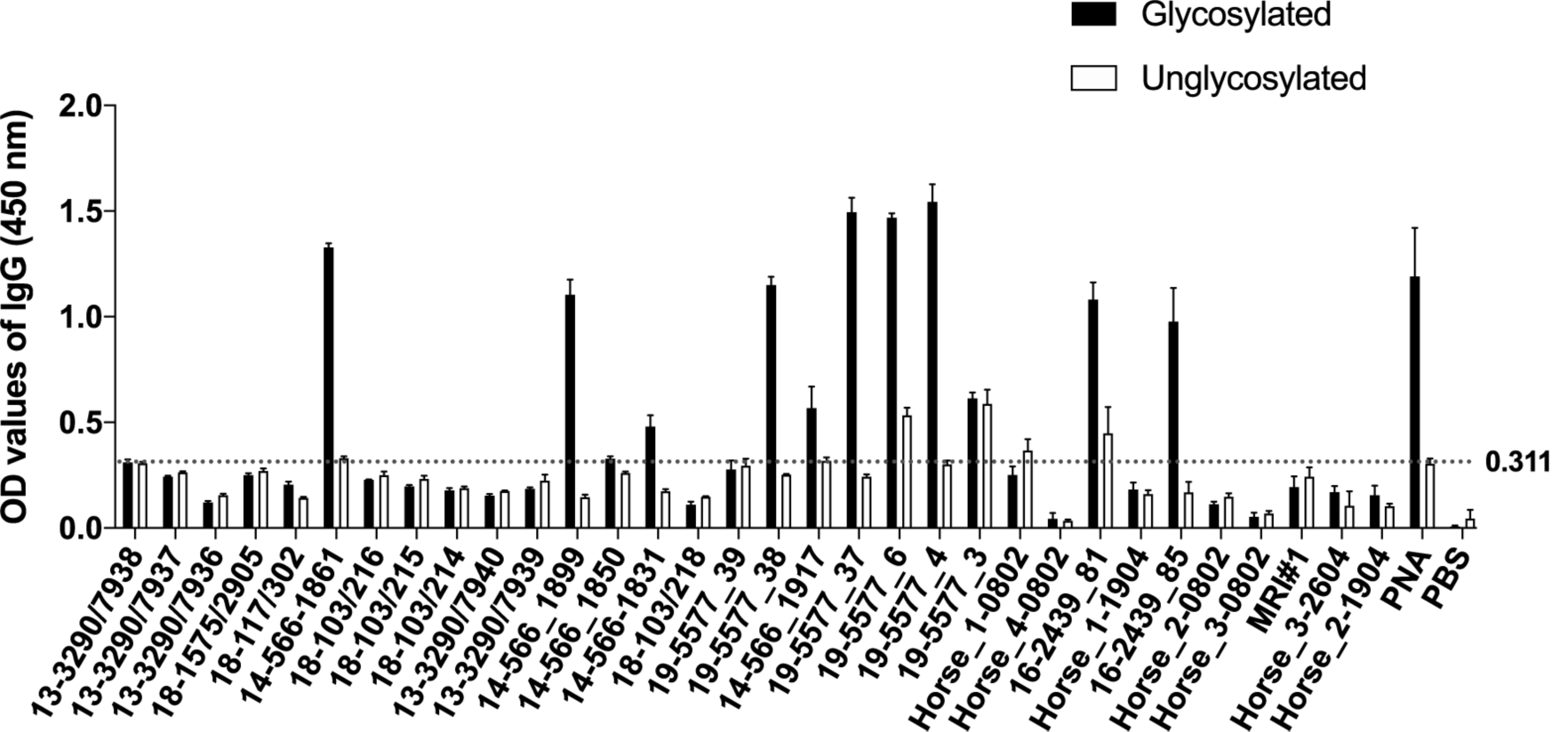
ELISA results using glycosylated and unglycosylated CtxB-BCAL2737a as a protein antigen. A panel of glanders positive and negative horse serum samples (see Table 1 for additional details) were analyzed by indirect ELISA (see Materials and Methods) using purified CtxB-BCAL2737a produced in *B. cenocepacia* K56-2 (glycosylated protein) and Δ*pglL* (unglycosylated negative control). Dotted line indicates the negative cutoff OD_450_ value for background color in the ELISA determined using 15 CFT negative serum samples. PNA (peanut agglutinin lectin) and PBS were also used as additional positive and negative controls, respectively. Three technical replicates were made for each sample.

To further validate the ELISA results, we carried out western blots using glycosylated CtxB-BCAL2737a and its unglycosylated form as a control. As expected, the results of western blot and ELISA agree, except for four samples (19-5577_3, 19-5577_38, Horse_2_(Poupée)_ D_19/04_, and Horse_3_(Princese)_ D_26/04_). The samples 19-5577_38 and 19-5577_3 were strongly positive in ELISA but negative in western blot, which could reflect differences in antibody titers related to the stage on disease at the time of serum extraction, since both sera correspond to infected animals in the same farm. In the other two samples, Horse_2_(Poupée)_ D_19/04_ and Horse_3_(Princese)_ D_26/04_, tested negative in ELISA but were positive in western blot (Fig. 5). These samples were considered true positives because they were also positive in both CFT and *B. mallei* ELISA. Moreover, same sensitivity and specificity was observed in western blot as compared to ELISA (Table 2), indicating that both methods can be used to detect anti-*O-*glycan antibodies in sera from infected horses.

**FIG 5.**
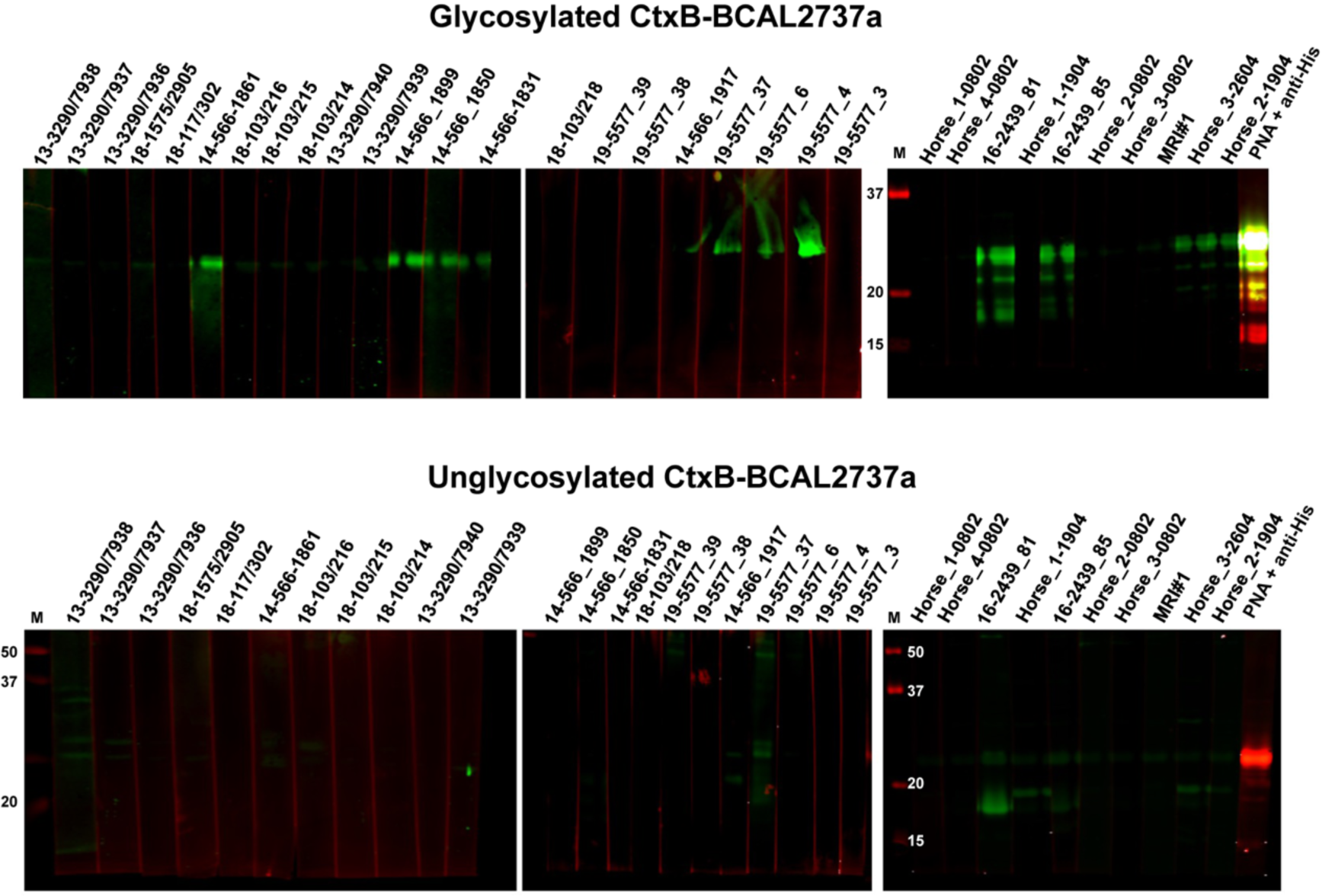
Western blot analysis of horse serum samples (see Table 1 for additional details) using glycosylated (top blots) and unglycosylated (bottom blots) CtxB-BCAL2737a. Blots were developed as described in Materials and Methods. As a control, glycosylated and unglycosylated CtxB-BCAL2737a were developed with PNA + anti-His (last lane in both series of blots). M, molecular mass markers; masses indicated in kilodaltons.

### Glycosylated CtxB-BCAL2737a can be used to detect anti-*O*-glycan antibodies in human serum samples from patients infected with various *Burkholderia* species

To examine whether glycosylated CtxB-BCAL2737a can also detect anti-*O*-glycan antibodies elicited from infections caused by other *Burkholderia* species, nine serum samples collected from patients infected with *B. multivorans, B. cenocepacia, B. mallei* and *B. pseudomallei* were used for western blots analysis. These samples had been reported to recognize an *O*-glycosylated DsbA1 protein from *Neisseria meningitidis* when expressed in *B. cenocepacia* (18). The results show that glycosylated CtxB-BCAL2737a but the not unglycosylated form can be recognized by antibodies present in these serum samples (Fig. 6), further emphasizing that glycosylated CtxB-BCAL2737a can be used to detect anti-*O*-glycan antibodies resulting from not only from *B. mallei* but also from other *Burkholderia*-associated infections.

**FIG 6.**
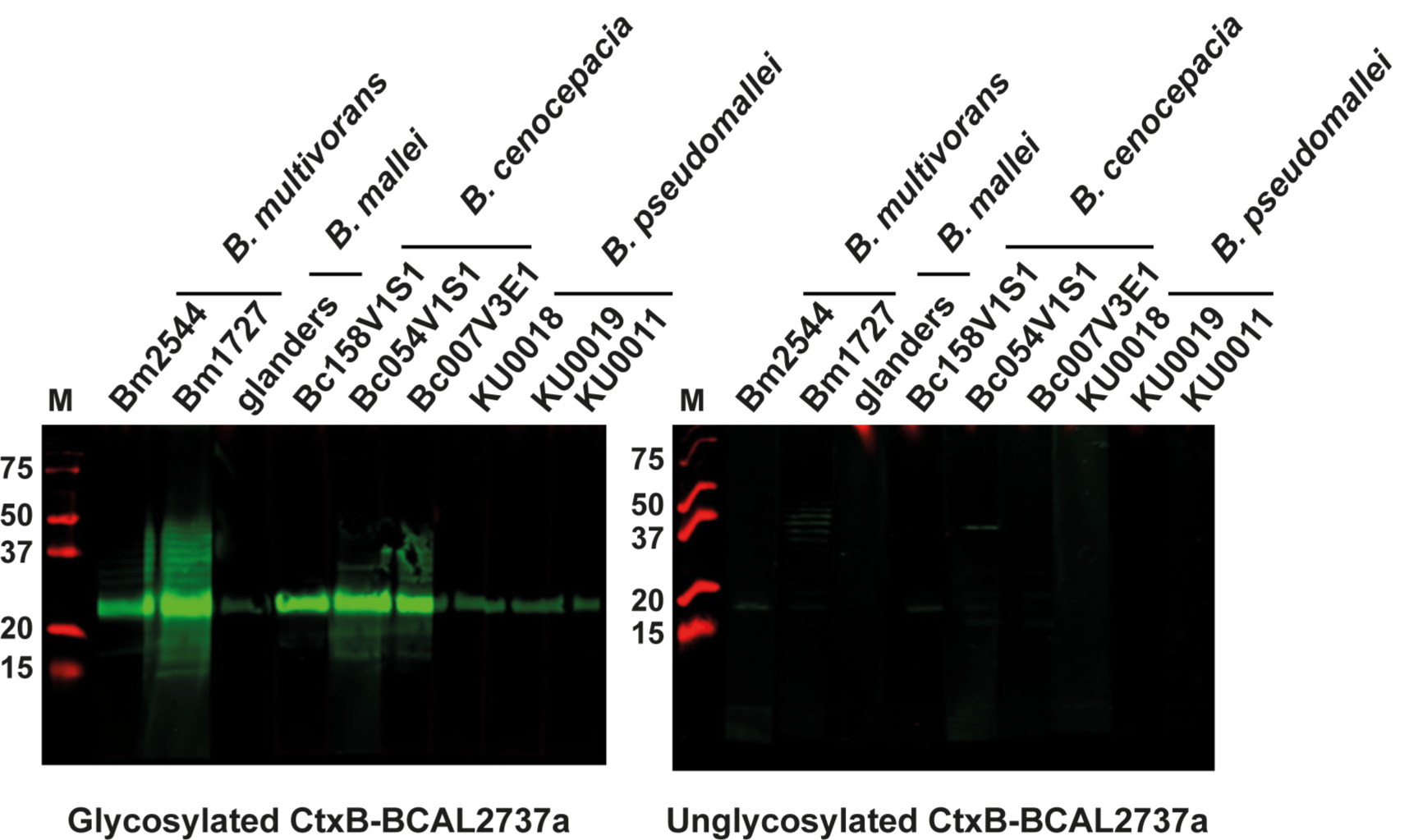
Western blot analysis of human serum samples from patients infected with the indicated *Burkholderia* species, as previously reported (20) using glycosylated and unglycosylated CtxB-BCAL2737a. Blots were developed as described in Materials and Methods. M, molecular mass markers; masses indicated in kilodaltons.

## DISCUSSION

Glanders poses a significant threat to both the equine industry and human health. Successful control of glanders infection requires reliable diagnostic tools. Although numerous methods (9-11, 13, 16, 18, 35) have been described for diagnosis of glanders, some have limitations (9-11, 18). Protein glycosylation, a post-translational modification that covalently attaches a glycan to a protein, is prevalent in Archaea and bacteria (36, 37). As some glycans contain unusual carbohydrates that are highly immunogenic, glycan-specific antibodies are attractive targets for diagnosis of bacterial infections and vaccine design (38). In this study, we exploited the conserved trisaccharide glycan synthesized by the *Burkholderia O*-glycosylation system to develop a glycoconjugate antigen for the diagnosis of glanders.

Previous studies on protein substrates for the *Burkholderia O*-glycosylation system revealed an abundant highly-glycosylated glycopeptide in *B. cenocepacia*, which could not be ascribed to any annotated gene (21). In this work, we identified and annotated this gene as BCAL2737a. The corresponding protein has 3 serine residues that can be *O-*glycosylated, as demonstrated by mass spectrometry analysis. An optimal *O*-glycosylation motif, WPAAA<u>S</u>AP (the underlined serine is the glycosylation acceptor site), required for recognition by the *Neisseria meningitidis* PglL *O*-Tase has been suggested (39), which is very similar to the sequences where we detect glycosylated serine residues in *Burkholderia* glycoproteins (20, 21, 32). The presence of three glycosylation sites in BCAL2737a, coupled to its small mass, make this protein an ideal carrier for presenting the trisaccharide. Our results demonstrated that *Burkholderia O*-glycosylation system can efficiently glycosylate the recombinant BCAL2737a protein with the correct trisaccharide structure. Moreover, fusion of this protein with the CtxB at N- and C-terminal locations did not affect the glycosylation of the BCAL2737a partner, suggesting that the glycosylation sites in BCAL2737a are accessible by the *Burkholderia* PglL *O*-Tase. This is consistent with the prediction that native glycosylation sites in bacterial proteins are located in flexible structures (40). However, we noticed that the native BCAL2737a polypeptide, as well as the BCAL2737a-CtxB fusion was unstable in the *O-*glycosylation-deficient mutants of *B. cenocepacia* K56-2. In bacterial *N-*glycosylation systems such as *Campylobacter jejuni*, loss of glycosylation affects the stability of the glycoproteome, which in turns reduces bacterial fitness and virulence (41, 42). Similar observations were made with protein *O*-glycosylation in *Burkholderia* (20), although the analysis of the proteome indicates that only some glycoproteins are highly reduced in abundance while others do not change significantly in the absence of glycosylation (32). Remarkably, BCAL2737a and BCAL2737a-CtxB were not detected in *O*-glycosylation defective mutants, indicating they are unstable in the absence of glycosylation. Our results demonstrated that unglycosylated BCAL2737a can only be stabilized when N-terminally fused with CtxB. The instability of BCAL2327a prevented us from using this protein in its unglycosylated form as a negative control in our serological assays, and to adsorb non-*O-* glycan potentially cross-reacting antibodies in tested sera. Together, our results suggest that *O-* glycosylation is required for the stability of BCAL2737a, possibly by assisting the folding of this polypeptide and preventing its degradation by periplasmic proteases. In this scenario, the N-terminally fused CtxB could help BCAL2737a to fold properly in the absence of glycosylation or at least adopt a proteolysis resistant conformation. Confirmation of this hypothesis awaits structural information of the chimeric protein.

Detection of anti-glycan antibodies using ELISA and western blot based on glycosylated CtxB-BCAL2737a suggest this antigen can be used for diagnosis of glanders, as high specificity (100%) and sensitivity (88.23%) was achieved by these two methods. Concerning the few false-negative results in both ELISA and western blot, several factors could affect the diagnostic reliability of ELISA and western blot, such as the purity and amount of the antigen, dilutions of test sample, and other interferences. We observed that the established protocol for ELISA worked well in detection of antibodies from samples of naturally-infected horses and did not have false positives in sera from noninfected animals. However, the ELISA failed to detect anti-*O*-glycan antibodies in samples obtained from an immunization trial with a heat-inactivated *B. mallei* preparation, especially from samples taken at 63 and 70 days after the trial’s initiation. In contrast, western blots were positive in later samples (see Table 1). It is likely that poor detection may be due to very low antibody titers against the *O-*glycan upon immunization with heat-inactivated *B. mallei* due to protein denaturation resulting in levels below the ELISA detection limit. This notion agrees with the finding that anti-*O-*glycan antibodies could be detectable by western blot upon increasing the amount of serum in the assay, suggesting that the heat-inactivated *B. mallei* antigen is a less potent inducer of anti-*O-*glycan antibodies compared to a natural infection. It is also possible that the major antibody isotype against the *O-*glycan in these sera was IgM, not IgG, due to the nature of the antigen and the immunization protocol. Detection of both IgM and IgG antibody classes in future trials may help improve the sensitivity of detection of anti-glycan antibodies in early infection. Further, our results showed that some samples appeared to have non-specific interactions with unglycosylated CtxB-BCAL2737a in ELISA. It is unlikely that these samples contain cross reactive antibodies that can recognize the carrier protein, because they did not interact with unglycosylated CtxB-BCAL2737a by western blot. Therefore, this might be attributed to contaminating proteins in the CtxB-BCAL2737a antigen preparation. Nevertheless, the combined use of ELISA and western blot was important to confirm inconclusive results. Further optimization and standardization of these assays are currently underway in our laboratory.

In conclusion, we have developed a recombinant glycoprotein antigen, CtxB-BCAL2737a, and have shown that ELISA and western blot assays based on the glycosylated CtxB-BCAL2737a can be used for diagnosis of glanders. Particularly, our ELISA test gave identical results compared to the semi-purified *B. mallei* ELISA (18). Combined with western blot, anti-*O-*glycan ELISA can add to the diagnostic tools for glanders. Diagnosing glanders, an OIE notifiable disease, have strong repercussions since it implies the slaughter of positive animals, and carries a negative impact on trade with other countries. Therefore, to have access to a battery of different tests with robust specificity is really important for a final decision. More broadly, our data support the notion that the conserved protein *O-*glycosylation system of *Burkholderia* can be used for diagnosis of other infections caused by these opportunistic bacteria.

## Acknowledgements

We thank Dr. Nora Madani for suppling equine sera from the experimental trial with *B. mallei* and Dr. Keren Turton for a critical review of the manuscript. We also than Dr. Pat Lenihan (Ireland), Dr. Vania Lucia Santana (Brazil), and Prof. Mohammed Sami A (Kuwait) for providing field negative and positive horse sera. This research was supported by a Medical Research Council Confidence in Concept grant (CD1617-CIC04) to M.A.V. and R.J.I. This work was supported by National Health and Medical Research Council of Australia (NHMRC) project grants awarded to NES (APP1100164). We thank the Melbourne Mass Spectrometry and Proteomics Facility of The Bio21 Molecular Science and Biotechnology Institute at The University of Melbourne for access to mass spectrometry infrastructure.

